# Host and infectivity prediction of Wuhan 2019 novel coronavirus using deep learning algorithm

**DOI:** 10.1101/2020.01.21.914044

**Authors:** Qian Guo, Mo Li, Chunhui Wang, Peihong Wang, Zhencheng Fang, Jie tan, Shufang Wu, Yonghong Xiao, Huaiqiu Zhu

**Author notes:** The authors contributed equally to this paper (Qian Guo, Mo Li, Chunhui Wang).

## Abstract

The recent outbreak of pneumonia in Wuhan, China caused by the 2019 Novel Coronavirus (2019-nCoV) emphasizes the importance of detecting novel viruses and predicting their risks of infecting people. In this report, we introduced the VHP (Virus Host Prediction) to predict the potential hosts of viruses using deep learning algorithm. Our prediction suggests that 2019-nCoV has close infectivity with other human coronaviruses, especially the severe acute respiratory syndrome coronavirus (SARS-CoV), Bat SARS-like Coronaviruses and the Middle East respiratory syndrome coronavirus (MERS-CoV). Based on our prediction, compared to the Coronaviruses infecting other vertebrates, bat coronaviruses are assigned with more similar infectivity patterns with 2019-nCoVs. Furthermore, by comparing the infectivity patterns of all viruses hosted on vertebrates, we found mink viruses show a closer infectivity pattern to 2019-nCov. These consequences of infectivity pattern analysis illustrate that bat and mink may be two candidate reservoirs of 2019-nCov.These results warn us to beware of 2019-nCoV and guide us to further explore the properties and reservoir of it.

**One Sentence Summary:** It is of great value to identify whether a newly discovered virus has the risk of infecting human. Guo *et al*. proposed a virus host prediction method based on deep learning to detect what kind of host a virus can infect with DNA sequence as input. Applied to the Wuhan 2019 Novel Coronavirus, our prediction demonstrated that several vertebrate-infectious coronaviruses have strong potential to infect human. This method will be helpful in future viral analysis and early prevention and control of viral pathogens.

## Main Text

The recently outbreak of pneumonia in Wuhan, China has brought the 2019 Novel Coronavirus (2019-nCoV) into our sight closely. The first case of unknown cause of pneumonia was found on 12 December 2019, and was later determined as a non-SARS novel coronavirus by Chinese Center for Disease Control and Prevention (China CDC). This virus has now caused a total of 217 confirmed human infections in China and three deaths, reported by Chinese media on 20 January 2020. Yet the virus has trend to spread out of China, since one case in Thailand, one case in Japan and two cases in Korea had been sequentially reported since 15 January 2020 *(1)*. Most of the patients are reported to have some link to seafood and animal market, which indicates the virus may spread from animals *(2)*. And the possibility of human-to-human transmission of the virus is still under investigation. The situation is severe and it is urgent for us to have a better understanding towards this virus for further prevention and control. Meanwhile, the outbreak of pneumonia caused by 2019-nCoV warns us the importance of predicting the risk of novel virus infection of human. At present, six full genomes of 2019-nCoV have been submitted on GISAID *(3)* by China CDC, *etc*., and among them one has been released on GenBank *(4)* (Accession: MN908947).

Currently there is urgent need to identify the hosts potentially infected by the 2019-nCoV viruses. Large amounts of infectious viruses can be obtained from domesticated animals, birds, vector animals and environments. These viruses may pose a great threat to human since lots of viruses that are safe in animals may potentially infect human. In addition, owing to high mutation rate in viruses, there are also non-human-infectious viruses that are originally spread in animals, but mutate to infect human later *(5)*. Besides, any unexpected hosts of novel viruses are also concerning.

To detect the potential host and pathogenicity of new viruses, the conventional way is based on their sequence similarity to known viruses, by either building phylogenetic tree or blast. However, this method seems to be rudimentary with several limitations. In fact, owing to the intrinsic heterogeneity within genome, different sequence fragments of the novel virus may be assigned to quite different known viruses with identical similarity by local alignment, which consequently leads to the ambiguous predictions of the hosts of novel viruses. In addition, the routine method, phylogenetic tree analysis, always utilizes the whole genomes assembled by shotgun reads, which is frequently influenced by the sequencing and assembly mistakes. Moreover, when the potential hosts of novel viruses were predicted from a phylogenetic perspective, people usually accepted the assumption that the viruses with closer phylogenetic relationships have common hosts, pathogenicity and infection behavior. However, this assumption has proved unsustainable, because the abilities of infecting human are varied even within genus and most viruses are harmless to human. For example, *Betacoronavirus*, a genus in *Coronavirinae*, containing both viruses that can infect human, such as SARS, and viruses that are not reported to infect human, such as *Rousettus* bat coronavirus HKU9.

Until now, there are several works on the identification of host of viruses, such as HostPhinder *(6)*, WIsH *(7)*, which all predict hosts for bacteriophages. However they were not designed for non-phage virus host-prediction, especially the virus causing Zoonotic diseases. To this end, we report the prediction results of the host of 2019-nCoV using the method, VHP (Virus Host Prediction), developed based on deep learning algorithm. The viral sequences data released before 2018 were used to build the training set, while those released after 2018 were utilized for testing. The dataset *(4)* for training and testing includes genomes of all DNA viruses, coding sequences of all RNA viruses and their host information in GenBank. In VHP’s prediction result of 2019-nCoV, while the values reflect the infectivity, the score pattern and p-value pattern reflect the infectivity pattern of the novel virus.

With the whole genome sequences released online, we predicted the potential hosts of 2019-nCoV, as well as other 44 Coronaviruses in NCBI refseq *(8)* and four Bat SARS-like Coronaviruses in GenBank *(4)*. It turns out that six genomes of 2019-nCoV all have a high possibility (*p-value*<0.05) to infect human (as shown in Table 1, and Table S2). Besides, most of the reported human-infective Coronaviruses *(9-14)* are assigned the lowest *p-values* predicted by the VHP method (Table S1). The similar probabilities of 2019-nCoV and other human Coronaviruses illustrates the high risk of 2019-nCoV. Part of the *p-value* results is shown in Table 1, which is ranked by the decreasing order of the *p-value* for a virus to infect human.

**Table 1.**
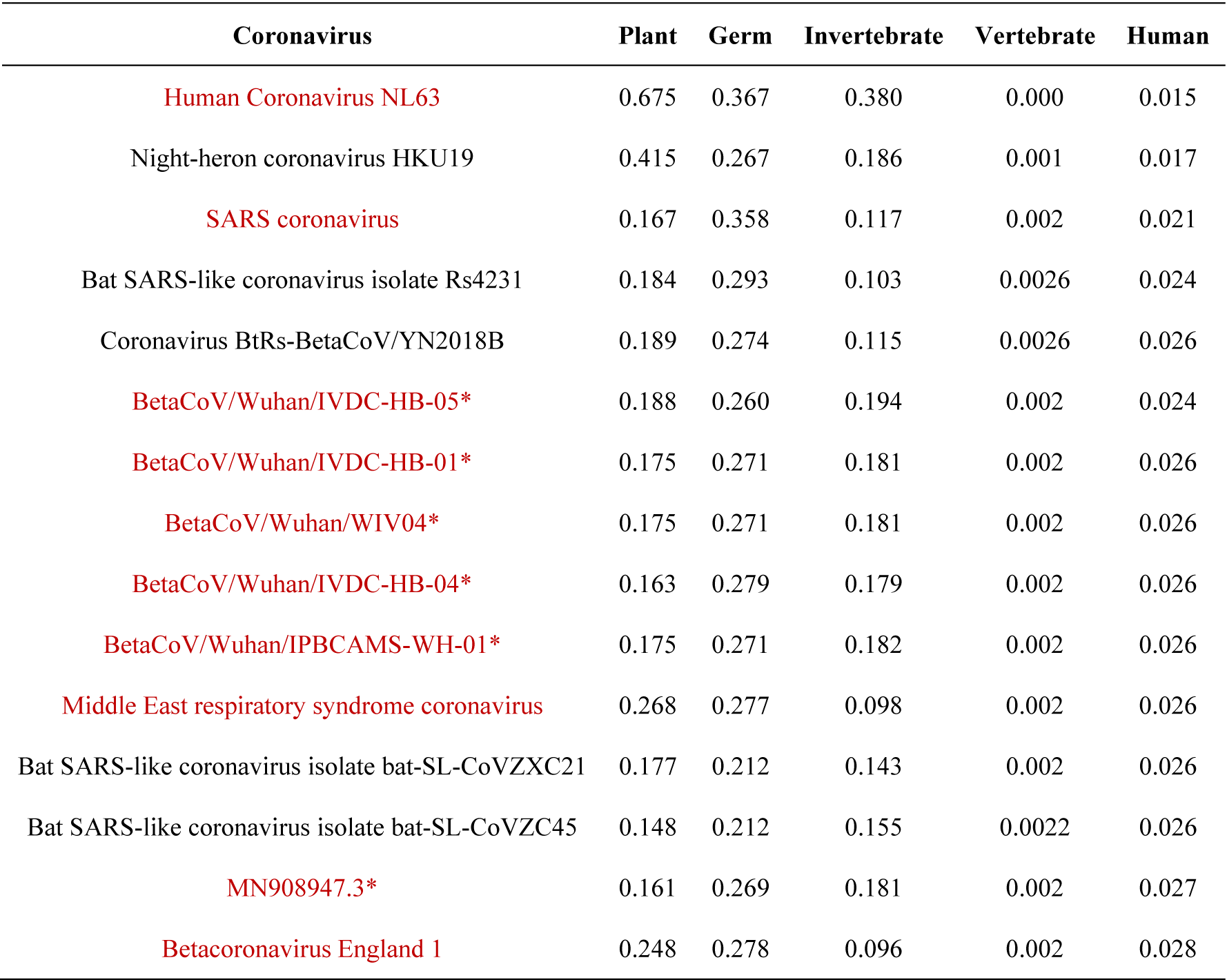
*P-values* of Potential Hosts of 50 Coronaviruses provided by the VHP method (human coronaviruses are colored red, 2019- nCoVs are marked with *).

The prediction results warn us that the 2019-nCoV have strong potential of infecting human similar to SARS, Bat SARS-like Coronaviruses and MERS. While SARS was assigned slightly lower *p-value* on “Human”, Bat SARS-like Coronaviruses were given closer *p-value* to 2019-nCoVs, which indicates that 2019-nCoV is probable to be less infectious within human than SARS but as infectious as Bat SARS-like Coronaviruses. Furthermore, we explore the reservoir of 2019-nCoVs with its infectivity pattern. As is shown in Table S3, compared with the *p-values* of other host types, “vertebrate” is given the least *p-value* for 2019-nCoVs predicted by VHP, which illustrates that the reservoir of 2019-nCoV may be vertebrate. Compared to the Coronaviruses infecting other vertebrates, Coronaviruses in bat are assigned with more similar infectivity patterns with 2019-nCoVs by VHP, which illustrates that bat is more likely to be the reservoir of 2019-nCoVs than other known vertebrate hosts of Coronaviruses. To explore other potential reservoirs, we scored all the available vertebrate viruses in GenBank, filtered viruses that have similar infectivity pattern with 2019-nCoV (Table S4). Results show that detailed hosts information of the filtered viruses include Canine, Porcine, Mink, Tortoise and Feline. By comparing the infectivity patterns of all viruses hosted on these vertebrates, we found Mink viruses show a closer infectivity pattern to 2019-nCov and thus we inferred Mink may be another candidate infection source of 2019-nCov.

To evaluate whether 2019-nCoVs are close to SARS, Bat SARS-like Coronaviruses and MERS genetically, we built a phylogenetic tree with the whole genome sequences of all the Coronaviruses utilized above, which showed that 2019-nCoVs have a closer phylogenetic relationship with SARS and Bat SARS-like Coronaviruses than other Coronaviruses (as shown in Fig. 1). In regard to protein level, we predicted the genes of 2019-nCoVs and annotated them and we found that the annotation results of all the six 2019-nCoV genomes are identical, including seven functional proteins (pp1a, E2 glycoprotein precursor, hypothetical protein sars3a, matrix protein, hypothetical protein sars6, hypothetical protein sars7a, nucleocapsid protein) verified in SARS-CoV-like coronavirus *(15)*. All these results illustrate the efficiency of our computational prediction and the high risk of 2019-nCoV and guide us to have a closer look at the similarity and differences of 2019-nCoV and SARS-CoV-like Coronaviruses.

**Fig. 1.**
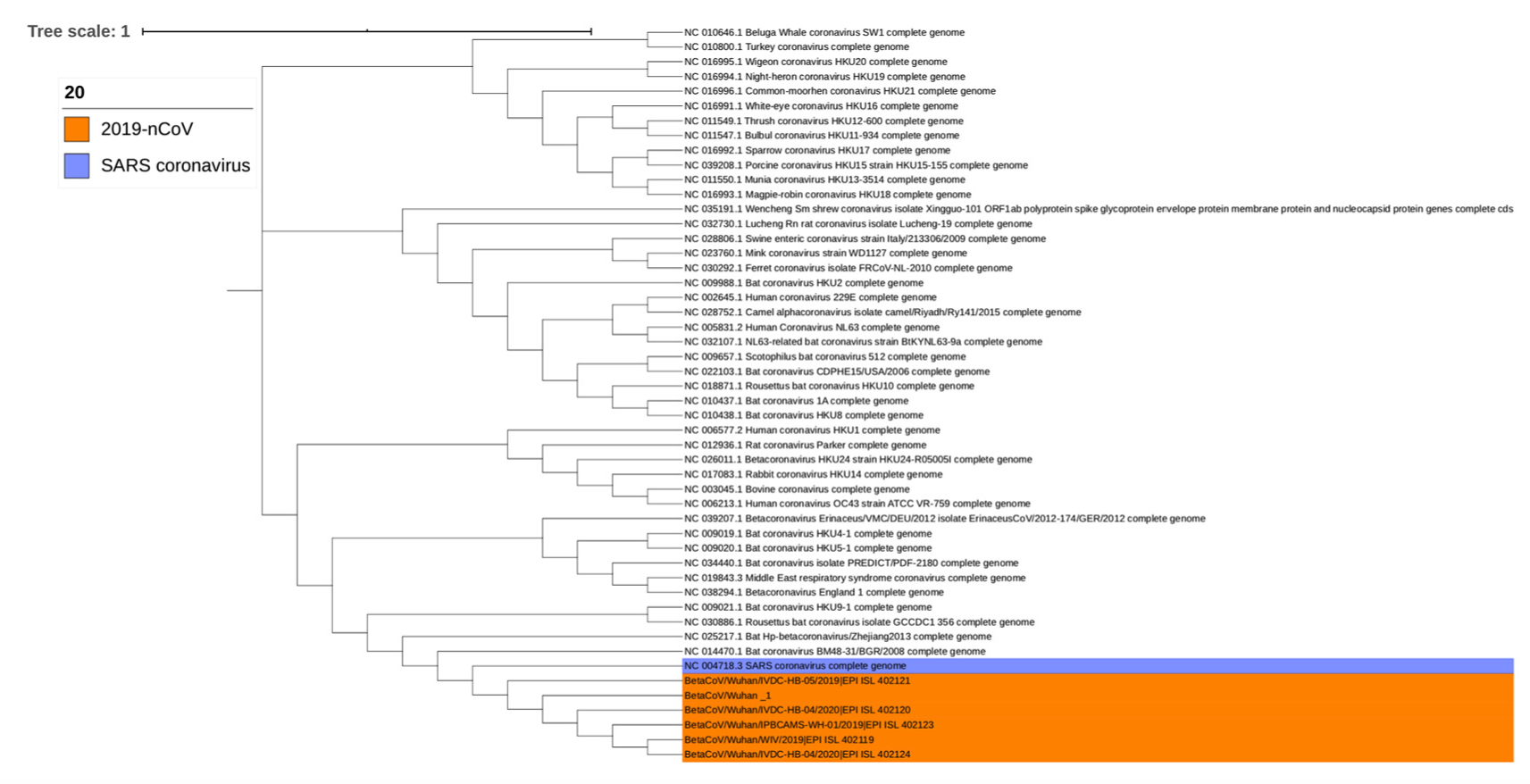
Phylogenetic tree of 2019-nCoV and 44 Coronaviruses.

To explore the differences of 2019-nCoVs, SARS and SARS-CoV-like Coronavirus, we then analyzed their core genes (more than 95% AAS identity) and nonsynonymous SNPs (single-nucleotide polymorphisms) (Table S3). We found that the six 2019-nCoVs are more similar to two Bat SARS-like Coronaviruses (MG772933 and MG772934) than SARS and other SARS-CoV-like Coronaviruses. In addition, the core gene SNP analysis shows that “BetaCoV/Wuhan/IVDC-HB-04” and “BetaCoV/Wuhan/IVDC-HB-05/2019” are divergent with the other four 2019-nCoVs, which means that there may be some mistakes in sequencing and assembly of these two viruses. By core gene and accessary gene analysis of the left four 2019-nCoVs and the two Bat SARS-like Coronaviruses, we found three core genes: two ORFs and one functional gene ymdB. While the four 2019-nCoVs have no SNPs on the two predicted ORFs, they have two nonsynonymous SNPs on ymdB, which indicates that these 2019-nCoV may derived from one origin. Besides, compared with the two Bat SARS-like Coronaviruses, 2019-nCoVs have 4, ∼130, ∼220 nonsynonymous SNPs on the three core genes respectively, which shows the differences among 2019-nCoV and SARS-CoV-like Coronaviruses.

Except for 2019-nCoVs, our predictions for all coronaviruses indicate that Night-heron coronavirus HKU19, which is reported to infect a vertebrate, Night-heron, also has the strong potential to infect human. Coronaviruses are a large family of viruses which usually circulate among animals which will cause fatal diseases, once they mutate and evolve to be able to infect human. For example, severe acute respiratory syndrome coronavirus (SARS-CoV) and Middle East respiratory syndrome coronavirus (MERS-CoV) are the most impressive ones. In November 2002, SARS-CoV found its first victim in Foshan, mainland China, and soon spread to all over the country by travelling with potential patients *(9)*. Different from other human-derived coronaviruses, SARS-CoV leads to severe acute respiratory syndrome (SARS), with a fatality rate of up to ∼10%. By July 2003, SARS-CoV had caused 8069 cases in 27 countries, 774 of them died *(16)*. Another distinctive coronavirus, known as MERS-CoV, was isolated from a man died of severe pneumonia and organ failure in Saudi Arabia at June 2012 *(10)*. MERS-CoV has infected 2,494 individuals in 27 countries since 2012, 858 of them died *(17)*. A much higher fatality rate (∼35%) has been shown in Middle East respiratory syndrome. Large epidemics of SARS and MERS have finished already, but follow-up studies never stop. High lethality and high human-to-human infectivity shown in these two diseases have raised alarm all over the world. Therefore, it’s urgent for us to seek out a new way and establish defensive lines for public health.

To sum up, in this report we predicted the potential of infecting human of 2019-nCoV, with implication of the risk of the 2019-nCoV. Aiming to give reliable predicted hosts and potential of infecting human of novel virus, VHP could play an important role in public health service and provide strong assistance for take precautions of novel viruses which have potential to infect human.

At the end of this report, we provide a brief description of VHP method and the verification of the algorithm. To construct the VHP model, we utilized a Bi-path Convolutional Neural Networks (BiPathCNN) *(18)*, where each viral sequence was represented by one-hot matrix of its base and codon separately. Considering difference in input sequence lengths, two BiPathCNNs (BiPathCNN-A and BiPathCNN-B) were built for predicting hosts of viral sequences from 100 bp to 400 bp and 400 bp to 800 bp respectively. The dataset *(7)* for training and testing includes genomes of all DNA viruses, coding sequences of all RNA viruses and their host information in GenBank. In order to develop a method expert in predicting the potential host types of novel viruses, the viral sequences data released before 2018 were used to build the training set, while those released after 2018 were utilized for testing. We integrated the hosts of viruses into five types, including plant, germ, invertebrate, vertebrate and human. The host subtypes contained in these five types are listed in Table 2 in detail. In the practical application for viral sequences, with the input of viral nucleotide sequence, VHP will output five scores for each host type, reflecting the infectivity within each host type respectively. Furthermore, VHP provides five *p-values*, statistical measures of how distinct the infections are compared with non-infection events. In order to evaluate the performance of VHP, we compare AUC (Area Under Curve) of both blast and VHP. The comparison result shows that VHP performs with evidently higher AUC in average (shown in Table 3).

**Table 2.**
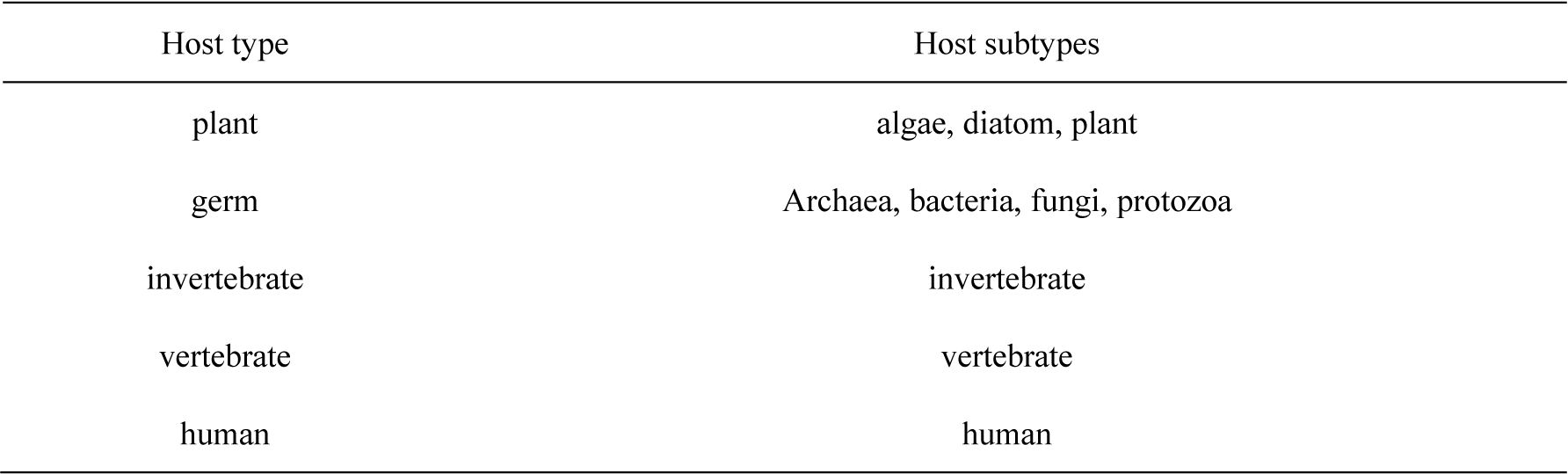
Host types and subtypes integrated in them.

| Host type | Host subtypes |
| --- | --- |
| plant | algae, diatom, plant |
| germ | Archaea, bacteria, fungi, protozoa |
| invertebrate | invertebrate |
| vertebrate | vertebrate |
| human | human |

**Table 3.** Comparison of AUC of blast and VPH

| Tool | Length of reads | Plant | Germ | Invertebrate | Vertebrate | Human |
| --- | --- | --- | --- | --- | --- | --- |
| VHP | 100-400bp | 0.860 | 0.948 | 0.924 | 0.927 | 0.972 |
|  | 400-800bp | 0.957 | 0.990 | 0.987 | 0.988 | 0.992 |
| Blast | 100-400bp | 0.616 | 0.747 | 0.798 | 0.739 | 0.830 |
|  | 400-800bp | 0.692 | 0.796 | 0.833 | 0.782 | 0.853 |

## Acknowledgments

We acknowledge China CDC; Wuhan Institute of Virology, Chinese Academy of Sciences; Institute of Pathogen Biology, Chinese Academy of Medical Sciences & Peking Union Medical College for genome sequencing sharing. We also acknowledge GISAID (https://www.gisaid.org/) for facilitating open data sharing. Part of the analysis was performed on the High Performance Computing Platform of the Center for Life Science of Peking University.

## Data and materials availability

All data is available in the main text or the supplementary materials.

## References

1. Centers for Disease Control and Prevention, 2019 Novel Coronavirus (2019-nCoV), Wuhan, China (2019); https://www.cdc.gov/coronavirus/2019-nCoV/summary.html.

2. J. Cohen, D. Normile, New SARS-like virus in China triggers alarm. Science. 367, 234–235 (2020).

3. GISAID, Newly discovered betacoronavirus, Wuhan 2019-2020 (2019); https://www.gisaid.org/.

4. J. R. Brister, D. Ako-Adjei, Y. Bao, O. Blinkova, NCBI viral genomes resource. Nucleic Acids Res. 43, D571–D577 (2015).

5. Controlprevention C F D, Swine-origin influenza A (H3N2) virus infection in two children--Indiana and Pennsylvania. Morb. Mortal. Wkly. Rep. 2011, 1213–1215 (2011).

6. J. Villarroel, K. A. Kleinheinz, V. I. Jurtz, H. Zschach, O. Lund, M. Nielsen, M. V. Larsen, HostPhinder: A Phage Host Prediction Tool. Viruses. 8, 10.3390/v8050116 (2016).

7. C. Galiez, M. Siebert, F. Enault, J. Vincent, J. Söding, WIsH: who is the host? Predicting prokaryotic hosts from metagenomic phage contigs. Bioinformatics. 33, 3113–3114 (2017).

8. E. W. Sayers, M. Cavanaugh, K. Clark, J. Ostell, K. D. Pruitt, I. Karsch-Mizrachi, GenBank. Nucleic Acids Res. 48, D84–86 (2020).

9. J. Ren, N. A. Ahlgren, Y. Y. Lu, J. A. Fuhrman, F. Sun, VirFinder: a novel k-mer based tool for identifying viral sequences from assembled metagenomic data. Microbiome. 5, 69 (2017).

10. N. S. Zhong, B. J. Zheng, Y. M. Li, Poon, Z. H. Xie, K. H. Chan, P. H. Li, S. Y. Tan, Q. Chang, J. P. Xie, X. Q. Liu, J. Xu, D. X. Li, K. Y. Yuen, Peiris, Y. Guan, Epidemiology and cause of severe acute respiratory syndrome (SARS) in Guangdong, People’s Republic of China, in February, 2003. Lancet. 362, 1353–1358 (2003).

11. A. M. Zaki, S. V. Boheemen, T. M. Bestebroer, A. D. Osterhaus, & R. A. Fouchier, Isolation of a novel coronavirus from a man with pneumonia in Saudi Arabia. N. Engl. J. Med. 367, 1814–1820 (2012).

12. M. A. Marra, S. J. Jones, C. R. Astell, R. A. Holt, A. Brooks-Wilson, Y. S. Butterfield, J. Khattra, J. K. Asano, S. A. Barber, S. Y. Chan, A. Cloutier, S. M. Coughlin, D. Freeman, N. Girn, O. L. Griffith, S. R. Leach, M. Mayo, H. McDonald, S. B. Montgomery, P. K. Pandoh, A. S. Petrescu, A. G. Robertson, J. E. Schein, A. Siddiqui, D. E. Smailus, J. M. Stott, G. S. Yang, F. Plummer, A. Andonov, H. Artsob, N. Bastien, K. Bernard, T. F. Booth, D. Bowness, M. Czub, M. Drebot, L. Fernando, R. Flick, M. Garbutt, M. Gray, A. Grolla, S. Jones, H. Feldmann, A. Meyers, A. Kabani, Y. Li, S. Normand, U. Stroher, G. A. Tipples, S. Tyler, R. Vogrig, D. Ward, B. Watson, R. C. Brunham, M. Krajden, M. Petric, D. M. Skowronski, C. Upton, R. L. Roper, The Genome sequence of the SARS-associated coronavirus. Science. 300, 1399–1404 (2003).

13. National Center for Biotechnology Information, Search database BtVs-BetaCoV/SC2013, complete genome, GenBank: KX285223.1; https://www.ncbi.nlm.nih.gov/nuccore/KJ473821.1.

14. L. V. D. Hoek, K. Pyrc, M. F. Jebbink, W. Vermeulen-Oost, R. J. Berkhout, K. C. Wolthers, P. M. W. Dillen, J. Kaandorp, J. Spaargaren, B. Berkhout, Identification of a new human coronavirus. Nat Med. 10, 368–373 (2004).

15. B. Buchfink, C. Xie, D. H. Huson. Fast and sensitive protein alignment using DIAMOND. Nat Methods. 12, 59–60 (2015).

16. World Health Organization, Summary of probably SARS cases with onset of illness from 1 November 2002 to 31 July 2003 (2003); http://www.who.int/csr/sars/country/table2004_04_21/en/.

17. World Health Organization. Middle East Respiratory Syndrome Coronavirus (MERS-CoV), MERS Monthly Summary, November 2019 (2019); http://www.who.int/emergencies/mers-cov/en/.

18. Z. Fang, J. Tan, S. Wu, M. Li, C. Xu, Z. Xie, H. Zhu, PPR-Meta: a tool for identifying phages and plasmids from metagenomic fragments using deep learning. Gigascience. 8, giz066 (2019).

19. D. C. Richter, F. Ott, A. F. Auch, R. Schmid, D. H. Huson, MetaSim: a sequencing simulator for genomics and metagenomics. PLoS One. 2008, 10.1371/journal.pone.0003373 (2008).

20. F. Sievers, A. Wilm, D. Dineen, T. J. Gibson, K. Karplus, W. Li, R. Lopez, H. McWilliam, M. Remmert, J. Söding, J. D. Thompson, D. G. Higgins, Fast, scalable generation of high-quality protein multiple sequence alignments using Clustal Omega. Mol Syst Biol. 7, 10.1038/msb.2011.75 (2011).

21. S. Kumar, G. Stecher, M. Li, C. Knyaz, K. Tamura, MEGA X: Molecular Evolutionary Genetics Analysis across Computing Platforms. Mol Biol Evol. 35, 1547–1549 (2018).

22. T. Seemann, Prokka: rapid prokaryotic genome annotation. Bioinformatics. 30, 2068–2069 (2014).

23. B. Buchfink, C. Xie, D. H. Huson. Fast and sensitive protein alignment using DIAMOND. Nat Methods. 12, 59–60 (2015).

24. A. J. Page, C. A. Cummins, M. Hunt, V. K. Wong, S. Reuter, M. T. Holden, M. Fookes, D. Falush, J. A. Keane, J. Parkhill, Roary: rapid large-scale prokaryote pan genome analysis. Bioinformatics. 31, 3691–3693 (2015).

25. Z. Yang, PAML: a program package for phylogenetic analysis by maximum likelihood. Comput Appl Biosci. 1997, 10.1093/bioinformatics/13.5.555 (1997).

