## Supplementary Materials for "Host and infectivity prediction of Wuhan 2019 novel coronavirus using deep learning algorithm"

### Datasets construction for training and testing

We downloaded all genomes of viruses which are annotated with host information from GenBank in July 9th, 2019. The viral sequences released before 2018 including 55283 genomes were used to build the training dataset, while those released after 2018 including 7756 genomes were utilized for testing. 1000,000 and 100,000 sequences were simulated by MetaSim*(19)* for training and testing respectively.

### Construction of VHP algorithm

<u>Mathematical representation of DNA sequences</u>

As the input of VHP, each DNA sequence will be converted to one-hot matrix of its base (BOH) and codon (COH) separately. In the coding of BOH, each consecutive base of a query sequence linked by its complementary strand is transformed with the base corresponding vectors: A: [0,0,0,1]; C: [0,0,1,0]; G: [0,1,0,0]; T: [1,0,0,0]. For COH, each query sequence is represented by the direct conjunction of its six phases (shown in figure S1) and then coded with the codon corresponding vectors with the length of 64 (identical to the number of codon type), similar to the base corresponding vectors.

Consequently, for a input sequence of length L, it will be transformed to a BOH matrix, with the size of $2L\times4$, and a COH matrix, with the size of $2L\times64$.

<u>Model description</u>

To build the framework of VHP, we introduced BiPathCNN (citiation), containing two CNN paths dig information from BOH matrix and COH matrix respectively. After independent convolution and pooling operations in the beginning, the two paths are combined by a concatenation layer. Following a normalization layer, five prediction scores will be provided by

five sub-paths, corresponding to the host type of plant, germ, invertebrate, vertebrate and human individually. The architecture of VHP is shown in Figure S2.
To improve the performance of VHP to predict virial sequences, we trained two models, BiPathCNN-A and BiPathCNN-B, which are receive the input sequences of length ranging from 100 to 400bp and 400 to 800bp separately.

Sequence prediction
In the practical application for viral sequences, which are longer than 800 bp, a cut window will move along the long sequence without overlapping to separate it into suitable fragments for the trained BiPathCNN-A and BiPathCNN-B. Finally, VPH will calculate the final score by weighting and summing the prediction scores acquired from BiPathCNN-A and BiPathCNN-B. For example, a 2000 bp query sequence will be separated into three consecutive fragments, corresponding to the first 800 bp, the middle 800 bp and the last 400 bp of the query sequence. Then VPH predicts calculates the weighted average of these fragments with the weights of 800/2000, 800/2000, and 400/2000 respectively.

p-value calculation
While the prediction scores given by VHP reflect the likelihood of each host type as the real host of the input virus, we calculate the *p-values* of each score, statistical measures of how distinct the host scores are compared with non-host scores. For example, if a input viral sequence is given a prediction score of 0.4 to infect human, we will compare 0.4 to the scores of non-human-infectious virus in our dataset and provide the *p-values* as an judgement basis. If the *p-values* is less than 0.05, we regard the input virus has significantly different infectivity compared with non-human-infectious virus and the potential pathogen of human.

**Sequence similarity-based analysis**

To explore the similarities and the differences between 2019-nCoV and other Coronaviruses, especially SARS Coronavirus and MERS Coronavirus, which are given closer prediction scores with 2019-nCoV by VHP, we did phylogenetic analysis and protein alignment.

Phylogenetic analysis
The Phylogenetic tree of all the Coronaviruses (44 Coronaviruse genomes downloaded from NCBI refseq *(8)* on Jan 15th, 2020, 5 2019-nCoV genomes downloaded from gsaid and 1 2019-nCoV genome downloaded from GenBank (Accession: MN908947.3) . Multiple sequence alignment was done by Clustal Omega *(20)*.Maximum likelihood tree was built by MEGA X *(21)*. The result (Fig. 1) shows that 2019-nCoV has closer evolutionary distance from SARS Coronavirus among all the Coronaviruses.

Protein annotation
In this section, we predicted proteins in 2019-nCoV with Prokka*(22)* at first. And then we built protein library with the proteins in 44 Coronaviruses and aligned the predicted proteins to the protein library by DIAMOND*(23)*. The annotation results of all the six 2019-nCoV genomes are identical, including 7 functional proteins(pp1a, E2 glycoprotein precursor, hypothetical protein sars3a, matrix protein, hypothetical protein sars6, hypothetical protein sars7a, nucleocapsid

protein) verified in SARS-CoV-like coronaviruses. The protein annotation result is shown in Table S2.

**SNP analysis**

With the sequence similarity-based analysis, we found 2019-nCoV were similar to the reported SARS-CoV-like coronaviruses, which guides us to explore the differences among them. In this part, we did core gene analysis for 6 2019-nCoVs and all the SARS-CoV-like coronaviruses in GenBank with Prokka and Roary*(22, 24)* and detected nonsynonymous SNPs with PAML*(25)*. As the differences of 2 2019-nCoVs ("BetaCoV/Wuhan/IVDC-HB-04" and "BetaCoV/Wuhan/IVDC-HB-05/2019") from the left 4, which was mentioned in the main text, we searched core genes and nonsynonymous SNPs of the 4 2019-nCoVs compared with the released reference genomes (MG772933 and MG772934). The nonsynonymous SNP results are shown in Table S3.

**Fig. S1.**

Converting the initial sequence to input sequence of BiPathCNNs by conjugating six phases of the sequence.

5'-ACGTTCAGCAT- 3'

↓

Phase1: ACG, TTC, AGC

Phase2: CGT, TCA, GCA

Phase3: GTT, CAG, CAT

Phase4: ATG, CTG, AAC

Phase5: TGC, TGA, ACG

Phase6: GCT, GAA, CGT

↓

Conjunction: ACG, TTC, AGC, CGT, TCA, GCA, GTT, CAG, CAT, ATG, CTG, AAC, TGC, TGA, ACG, GCT, GAA, CGT

**Fig. S2.**

Construction of BiPathCNNs.

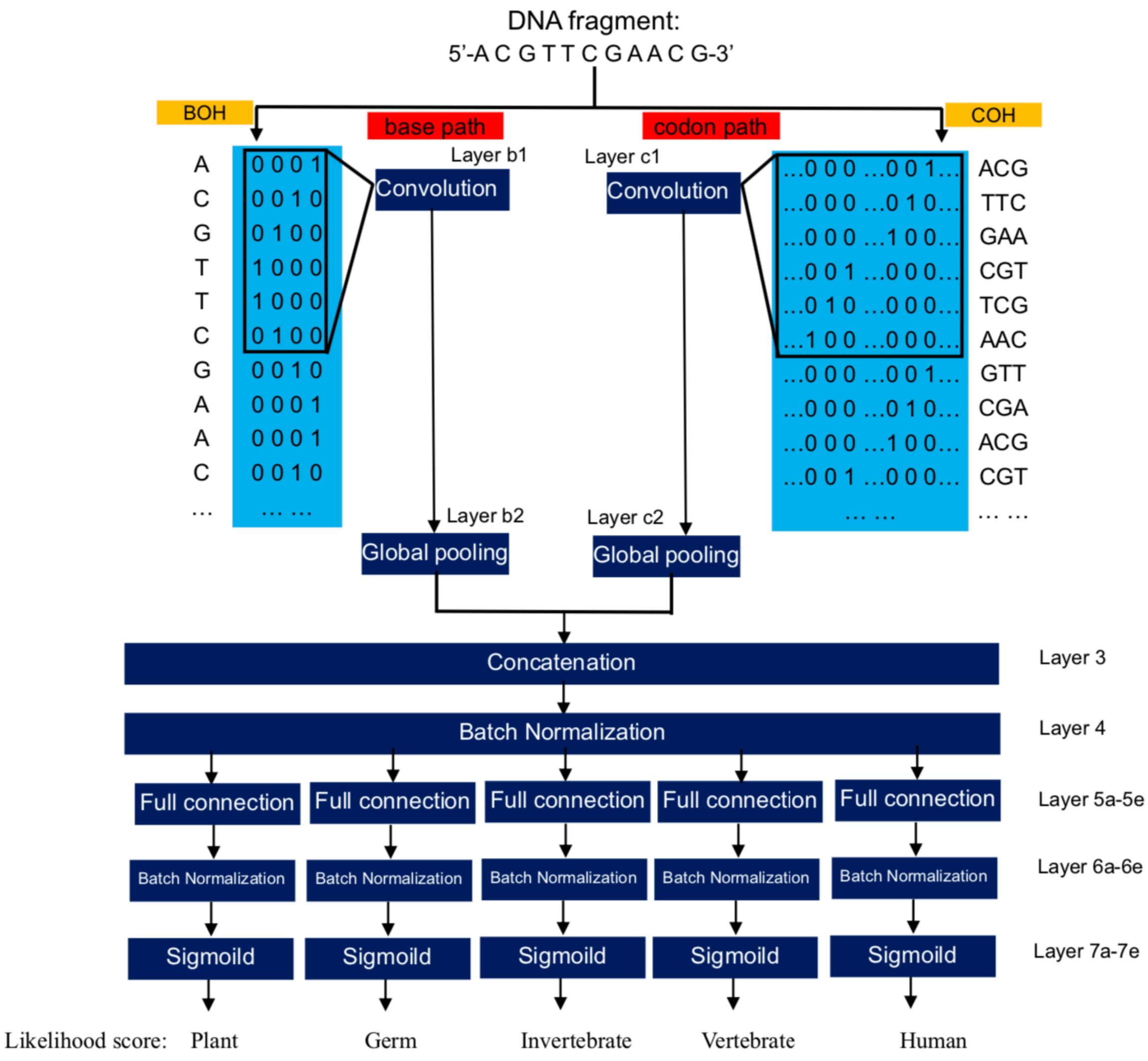

**Table S1.**

*P-values* of Potential Hosts of 54 Coronaviruses provided by VHP (human Coronaviruses are colored red and 2019-nCoVs are marked with *).

| Coronavirus | Plant | Germ | Invertebrate | Vertebrate | Human |
|---|---|---|---|---|---|
| Human Coronavirus NL63 | 0.675 | 0.367 | 0.380 | 0.000 | 0.015 |
| Night-heron coronavirus HKU19 | 0.415 | 0.267 | 0.186 | 0.001 | 0.017 |
| SARS coronavirus | 0.167 | 0.358 | 0.117 | 0.002 | 0.021 |
| Bat SARS-like coronavirus isolate Rs4231 | 0.184 | 0.293 | 0.103 | 0.0026 | 0.024 |
| Coronavirus BtRs-BetaCoV/YN2018B | 0.189 | 0.274 | 0.115 | 0.0026 | 0.026 |
| BetaCoV/Wuhan/IVDC-HB-05* | 0.188 | 0.260 | 0.194 | 0.002 | 0.024 |
| BetaCoV/Wuhan/IVDC-HB-01* | 0.175 | 0.271 | 0.181 | 0.002 | 0.026 |
| BetaCoV/Wuhan/WIV04* | 0.175 | 0.271 | 0.181 | 0.002 | 0.026 |
| BetaCoV/Wuhan/IVDC-HB-04* | 0.163 | 0.279 | 0.179 | 0.002 | 0.026 |
| BetaCoV/Wuhan/IPBCAMS-WH-01* | 0.175 | 0.271 | 0.182 | 0.002 | 0.026 |
| Middle East respiratory syndrome coronavirus | 0.268 | 0.277 | 0.098 | 0.002 | 0.026 |
| Bat SARS-like coronavirus isolate bat-SL-CoVZXC21 | 0.177 | 0.212 | 0.143 | 0.002 | 0.026 |
| Bat SARS-like coronavirus isolate bat-SL-CoVZC45 | 0.148 | 0.212 | 0.155 | 0.0022 | 0.026 |
| MN908947.3* | 0.161 | 0.269 | 0.181 | 0.002 | 0.027 |
| Betacoronavirus England 1 | 0.248 | 0.278 | 0.096 | 0.002 | 0.028 |
| Bat coronavirus isolate PREDICT | 0.220 | 0.320 | 0.097 | 0.002 | 0.028 |
| Thrush coronavirus HKU12-600 | 0.315 | 0.373 | 0.097 | 0.001 | 0.028 |
| Camel alphacoronavirus isolate camel | 0.318 | 0.366 | 0.355 | 0.000 | 0.029 |
| White-eye coronavirus HKU16 | 0.346 | 0.323 | 0.122 | 0.001 | 0.030 |
| Bulbul coronavirus HKU11-934 | 0.352 | 0.333 | 0.123 | 0.001 | 0.030 |
| Bat coronavirus BM48-31 | 0.221 | 0.378 | 0.229 | 0.001 | 0.031 |
| Human coronavirus 229E | 0.276 | 0.393 | 0.413 | 0.000 | 0.032 |
| Common-moorhen coronavirus HKU21 | 0.584 | 0.201 | 0.068 | 0.002 | 0.033 |
| Wigeon coronavirus HKU20 | 0.359 | 0.376 | 0.115 | 0.001 | 0.033 |
| Betacoronavirus Erinaceus | 0.435 | 0.380 | 0.139 | 0.001 | 0.033 |
| Human coronavirus OC43 strain ATCC VR-759 | 0.383 | 0.330 | 0.184 | 0.001 | 0.034 |
| Turkey coronavirus | 0.238 | 0.296 | 0.167 | 0.001 | 0.034 |
| Rousettus bat coronavirus HKU10 | 0.310 | 0.452 | 0.470 | 0.000 | 0.035 |

| | | | | | |
|---|---|---|---|---|---|
| Bovine coronavirus | 0.371 | 0.337 | 0.190 | 0.001 | 0.035 |
| Rabbit coronavirus HKU14 | 0.346 | 0.378 | 0.121 | 0.001 | 0.036 |
| Rat coronavirus Parker | 0.194 | 0.347 | 0.091 | 0.002 | 0.037 |
| Swine enteric coronavirus strain | 0.204 | 0.323 | 0.250 | 0.001 | 0.037 |
| Bat Hp-betacoronavirus | 0.175 | 0.361 | 0.110 | 0.002 | 0.037 |
| Munia coronavirus HKU13-3514 | 0.349 | 0.363 | 0.129 | 0.001 | 0.038 |
| Bat coronavirus HKU4-1 | 0.361 | 0.200 | 0.142 | 0.001 | 0.039 |
| Mink coronavirus strain WD1127 | 0.529 | 0.330 | 0.468 | 0.000 | 0.039 |
| Betacoronavirus HKU24 strain HKU24-R05005I | 0.210 | 0.378 | 0.066 | 0.002 | 0.040 |
| Ferret coronavirus isolate FRCoV-NL-2010 | 0.405 | 0.411 | 0.379 | 0.000 | 0.040 |

**Table S2.**

Results of protein annotation of 2019-nCovs. Proteins in the header represents 7 proteins in SARS. '+' represents having a blast hit.

| 2019-nCoV | Number of predicted proteins | Orf1a polyprotein (pp1a) | E2 glycoprotein precursor | Hypothetical protein sars3a | Matrix protein | Hypothetical protein sars6 | Hypothetical protein sars7a | Nucleocapsid protein |
|---|---|---|---|---|---|---|---|---|
| BetaCoV/Wuhan/IVDC-HB-01 | 9 | + | + | + | + | + | + | + |
| BetaCoV/Wuhan/IVDC-HB-04 | 15 | + | + | + | + | + | + | + |
| BetaCoV/Wuhan/IVDC-HB-05 | 10 | + | + | + | + | + | + | + |
| BetaCoV/Wuhan/IPBCAMS-WH-01 | 9 | + | + | + | + | + | + | + |
| BetaCoV/Wuhan/WIV04 | 9 | + | + | + | + | + | + | + |
| BetaCoV/Wuhan _1 | 9 | + | + | + | + | + | + | + |

**Table S3.**

The number of SNPs on 3 core genes of 6 2019-nCoVs and 2 SARS-CoV-like Coronaviruses.

| Virus pairs | Core gene1 | Core gene2 | Core gene3(gmdB) |
|---|---|---|---|
| IVDC-HB-01/ IPBCAMS-WH-01 | 0 | 0 | 2 |
| IVDC-HB-01/WIV04 | 0 | 0 | 0 |
| IVDC-HB-01/MN908947.3 | 0 | 0 | 0 |
| IVDC-HB-01/MG772933 | 4 | 125 | 207 |
| IVDC-HB-01/MG772934 | 4 | 130 | 220 |
| IPBCAMS-WH-01/WIV04 | 0 | 0 | 2 |
| IPBCAMS-WH-01/MN908947.3 | 0 | 0 | 2 |
| IPBCAMS-WH-01/MG772933 | 4 | 125 | 209 |
| IPBCAMS-WH-01/MG772934 | 4 | 130 | 223 |
| WIV04/ MN908947.3 | 0 | 0 | 0 |
| WIV04/MG772933 | 4 | 125 | 207 |
| WIV04/MG772934 | 4 | 130 | 220 |
| MN908947.3/MG772933 | 4 | 125 | 207 |
| MN908947.3/MG772934 | 4 | 139 | 220 |
| MG772933/MG772934 | 0 | 30 | 94 |

**Table S4.**

VHP's predictions for vertebrate viruses with close p-values with 2019-nCovs

| Vertebrate virus | Plant | Germ | Invertebrate | Vertebrate | Human |
|---|---|---|---|---|---|
| MG279127.1 Canine circovirus isolate 180 | 0.170 | 0.163 | 0.157 | 0.0019 | 0.027 |
| NC_038537.1 Bocavirus pig/SX/China/2010 NS1 (NS1), NP1 (NP1), VP1 (VP1), and VP2 (VP2) genes | 0.187 | 0.112 | 0.263 | 0.0021 | 0.026 |
| NC_023020.1 Porcine parvovirus 5 isolate IA469 clone 1 | 0.173 | 0.245 | 0.111 | 0.0021 | 0.026 |
| AY780926.1 Infectious pancreatic necrosis virus strain 6B1a segment B | 0.273 | 0.285 | 0.004 | 0.0021 | 0.027 |
| MG279120.1 Canine circovirus isolate 198 | 0.174 | 0.165 | 0.133 | 0.0021 | 0.025 |
| NC_020499.1 Canine bocavirus 1 isolate Con-161 | 0.171 | 0.248 | 0.065 | 0.0022 | 0.027 |
| MG001454.1 Mink circovirus strain HL24 | 0.169 | 0.228 | 0.151 | 0.0022 | 0.027 |
| NC_022104.1 Porcine partetravirus strain FMV10-1437266 | 0.206 | 0.269 | 0.084 | 0.0022 | 0.027 |
| NC_001916.1 Infectious pancreatic necrosis virus segment B | 0.336 | 0.216 | 0.003 | 0.0024 | 0.028 |
| NC_025890.1 Tortoise picornavirus strain 14-04 | 0.441 | 0.252 | 0.035 | 0.0026 | 0.025 |
| LT898442.1 Pasivirus A isolate SPaV-A_GER_L00721_2014 genome assembly | 0.058 | 0.374 | 0.128 | 0.0026 | 0.026 |
| NC_039044.1 Feline bocaparvovirus 3 isolate FBD1 | 0.182 | 0.288 | 0.054 | 0.0028 | 0.027 |

### Acknowledge table

We gratefully acknowledge the authors, originating and submitting laboratories of the sequences from GISAID's EpiFlu™ Database on which this research is based. The list is detailed below.

All submitters of data may be contacted directly via the GISAID (3) website www.gisaid.org

| Accession ID | Virus name | Location | Collection date | Originating lab | Submitting lab | Authors |
|---|---|---|---|---|---|---|
| EPI_ISL_402120 | BetaCoV/Wuhan/IVDC-HB-04/2020 | China/ Hubei Province / Wuhan City | 2020-01-01 | National Institute for Viral Disease Control and Prevention, China CDC | National Institute for Viral Disease Control and Prevention, China CDC | Wenjie Tan，Xiang Zhao，Wenling Wang，Xuejun Ma，Yongzhong Jiang，Roujian Lu，Ji Wang，Weimin Zhou，Peihua Niu，Peipei Liu，Faxian Zhan，Weifeng Shi，Baoying Huang，Jun Liu，Li Zhao，Yao Meng，Xiaozhou He，Fei Ye，Na Zhu，Yang Li，Jing Chen，Wenbo Xu，George F. Gao，Guizhen Wu |
| EPI_ISL_402119 | BetaCoV/Wuhan/IVDC-HB-01/2019 | China / Hubei Province / Wuhan City | 2019-12-30 | National Institute for Viral Disease Control and Prevention, China CDC | National Institute for Viral Disease Control and Prevention, China CDC | Wenjie Tan，Xiang Zhao，Wenling Wang，Xuejun Ma，Yongzhong Jiang，Roujian Lu, Ji Wang, Weimin Zhou，Peihua Niu，Peipei Liu，Faxian Zhan，Weifeng Shi，Baoying Huang，Jun Liu，Li Zhao，Yao Meng，Xiaozhou He，Fei Ye，Na Zhu，Yang Li，Jing Chen，Wenbo Xu，George F. Gao，Guizhen Wu |
| EPI_ISL_402121 | BetaCoV/Wuhan/IVDC-HB-05/2019 | China / Hubei Province / Wuhan City | 2019-12-30 | National Institute for Viral Disease Control and Prevention, China CDC | National Institute for Viral Disease Control and Prevention, China CDC | Wenjie Tan，Xuejun Ma，Xiang Zhao，Wenling Wang，Yongzhong Jiang，Roujian Lu，Ji Wang，Peihua Niu, Weimin Zhou, Faxian Zhan，Weifeng Shi，Baoying Huang，Jun Liu，Li Zhao，Yao Meng，Fei Ye，Na Zhu, Xiaozhou He，Peipei Liu, Yang Li，Jing Chen，Wenbo Xu，George F. Gao，Guizhen Wu |
| EPI_ISL_402124 | BetaCoV/Wuhan/WIV04/2019 | China / Hubei Province / Wuhan City | 2019-12-30 | Wuhan Jinyintan Hospital | Wuhan Institute of Virology, Chinese Academy of Sciences | Peng Zhou, Xing-Lou Yang, Ding-Yu Zhang, Lei Zhang, Yan Zhu, Hao-Rui Si, Zhengli Shi |
| EPI_ISL_402123 | BetaCoV/Wuhan/IPBCAMS-WH-01/2019 | China / Hubei Province / Wuhan City | 2019-12-24 | Institute of Pathogen Biology, Chinese Academy of Medical Sciences & Peking Union Medical College | Institute of Pathogen Biology, Chinese Academy of Medical Sciences & Peking Union Medical College | Lili Ren, Jianwei Wang, Qi Jin, Zichun Xiang, Yongjun Li, Zhiqiang Wu, Chao Wu, Yiwei Liu |
| EPI_ISL_402125 | BetaCoV/Wuhan-Hu-1/2019 | China | 2019-12 | unknown | National Institute for Communicable Disease Control and Prevention (ICDC) Chinese Center for Disease Control and Prevention (China CDC) | Zhang,Y.-Z., Wu,F., Chen,Y.-M., Pei,Y.-Y., Xu,L., Wang,W., Zhao,S., Yu,B., Hu,Y., Tao,Z.-W., Song,Z.-G., Tian,J.-H., Zhang,Y.-L., Liu,Y., Zheng,J.-J., Dai,F.-H., Wang,Q.-M., She,J.-L. and Zhu,T.-Y. |
